# Direct PCR offers a fast and reliable alternative to conventional DNA isolation methods for animal gut microbiomes

**DOI:** 10.1101/193664

**Authors:** Elin Videvall, Maria Strandh, Anel Engelbrecht, Schalk Cloete, Charlie K. Cornwallis

## Abstract

The gut microbiome of animals is emerging as an important factor influencing ecological and evolutionary processes. A major bottleneck in obtaining microbiome data from large numbers of samples is the time-consuming laboratory procedures, specifically the isolation of DNA and generation of amplicon libraries. Recently, direct PCR kits have been developed that circumvent conventional DNA extraction steps, thereby streamlining the laboratory process by reducing preparation time and costs. However, the reliability and efficacy of the direct PCR method for measuring host microbiomes has not yet been investigated other than in humans with 454-sequencing. Here, we conduct a comprehensive evaluation of the microbial communities obtained with direct PCR and the widely used MoBio PowerSoil DNA extraction kit in five distinct gut sample types (ileum – caecum – colon – faeces – cloaca) from 20 juvenile ostriches, using 16S rRNA Illumina MiSeq sequencing. We found that direct PCR was highly comparable over a range of measures to the DNA extraction method in caecal, colon, and faecal samples. However, the two methods recovered significantly different microbiomes in cloacal, and especially ileal samples. We also sequenced 100 replicate sample pairs to evaluate repeatability during both extraction and PCR stages, and found that both methods were highly consistent for caecal, colon, and faecal samples (r_s_ > 0.7), but had low repeatability for cloacal (r_s_ = 0.39) and ileal (r_s_ = −0.24) samples. This study indicates that direct PCR provides a fast, cheap, and reliable alternative to conventional DNA extraction methods for retrieving 16S data, which will aid future gut microbiome studies of animals.

## Introduction

It is becoming increasingly evident that the microbes animals harbour play an important role in regulating physiology and behaviour of individuals (McFall-Ngai *et al.* 2013). For example, the human gut contains around 39 trillion bacteria (Sender *et al.* 2016), which, together with other microorganisms, have large effects on human health and disease (Collins *et al.* 2012; Tremaroli & BÄckhed 2012; Gensollen *et al.* 2016). Although the varied and prominent effects of the gut microbiome on hosts has been brought to the forefront by studies on humans, research on non-human organisms is rapidly expanding and illustrating the importance of host microbiomes for a range of ecological and evolutionary processes (e.g. Ezenwa *et al.* 2012; Dill-Mcfarland *et al.* 2014; Waite & Taylor 2014; Tung *et al.* 2015; Knutie *et al.* 2017; Small *et al.* 2017; Groussin *et al.* 2017). This has been greatly aided by progress in sequencing technologies and genetic markers, such as 16S rRNA, that allow large numbers of bacterial communities to be characterised. However, the scope of microbiome studies continues to be limited by time-consuming laboratory procedures, in particular the isolation of DNA and the generation of amplicon libraries. As phenotypic variation is widespread in natural populations, the success of ecological and evolutionary studies on host microbiomes relies on large sample sizes, and so it is important to find fast, cost effective, and reliable ways of processing microbiome samples.

The conventional way of generating amplicon libraries for microbiome studies is to first extract and purify DNA, for example, using kits such as the MoBio PowerSoil DNA isolation kit. This procedure is recommended by the Earth Microbiome Project (Caporaso *et al.* 2012; Gilbert *et al.* 2014), and is widely used in human and non-human animal microbiota studies. The DNA extraction protocol involves mechanical and chemical lysis of cells, and a DNA purification procedure, which adds up to 32 separate steps (Table 1). One potentially faster and cheaper technique to prepare amplicon libraries is the recently developed direct PCR method. This method minimises the DNA isolation steps because DNA is simply extracted in a buffer with a 10 min 95 °C treatment prior to PCR amplification. To our knowledge, the accuracy of direct PCR versus DNA extraction for 16S sequencing of microbiomes has only been assessed once before, using human samples from four individuals, and with the discontinued 454 pyrosequencing technique (Flores *et al.* 2012). The results of that comparison suggested that the direct PCR method was a highly viable alternative to the DNA extraction method (Flores *et al.* 2012). This raises the question of whether direct PCR provides a cheaper and faster alternative to the conventional methods currently being used in non-human animal microbiome studies.

**Table 1.** Comparison of practical factors between DNA extraction and direct PCR methods for gut microbiome studies.

| Method | DNA extraction<sup>a</sup> | Direct PCR<sup>b</sup> |
| --- | --- | --- |
| Extraction time (h)<sup>c</sup> | ~8 h (1–2 days) | 45 min |
| Number of protocol steps | 32 | 3 |
| Kit cost/sample (USD) | 6.90<sup>d</sup> | 2.30<sup>e</sup> |
| Storage of extracted DNA | Long-term | Short-term |

<sup>a</sup>Information is given for DNA extraction with the PowerSoil-htp 96 Well Soil DNA Kit, 384 extractions (MO BIO Laboratories).
<sup>b</sup>Information is given for direct PCR with the Extract-N-Amp Plant PCR Kit, 100 extractions (Sigma-Aldrich).
<sup>c</sup>Time is estimated for parallel extraction of 96–192 samples.
<sup>d</sup>Price includes PCR reagents (Phusion High-Fidelity PCR Master Mix with HF Buffer, ThermoScientific) that are not part of the DNA extraction kit.
<sup>e</sup>Price includes PCR reagents that are part of the kit.

We therefore evaluated microbial communities obtained using the direct PCR approach (modified from Flores *et al.* 2012) compared to the conventional DNA extraction technique (MoBio PowerSoil DNA isolation kit) using 16S rRNA Illumina MiSeq sequencing. We examined the performance of these two techniques across the length of the gut (ileum – caecum – colon) and in two commonly used sample types for animal microbiome studies (faeces and cloacal swabs) of juvenile ostriches as a case study, given these samples are known to differ markedly in microbial composition (Videvall *et al.* 2017). Specifically, our aims were to test if direct PCR and DNA extraction methods differ in: 1) community diversity and distance, 2) specific bacterial taxa and abundances, and 3) repeatability of replicates from the same samples.

## Materials and methods

### Sample collection

Information of most materials and methods, including sample collection, has been outlined in detail by Videvall *et al.* (2017). Briefly, samples were collected from juvenile ostriches, *Struthio camelus*, kept at the Western Cape Department of Agriculture ostrich research facility in Oudtshoorn, South Africa. Samples were collected from 20 randomly selected birds (10 individuals four weeks old and 10 individuals six weeks old). Five different sample types were collected from each individual: faeces, cloacal swabs, and gut contents from the ileum, caecum, and colon, together with control swabs (Videvall *et al.* 2017). All procedures were approved by the Departmental Ethics Committee for Research on Animals (DECRA) of the Western Cape Department of Agriculture, reference number R13/90. The samples were collected between October 28 and November 12, 2014, in 2 ml plastic micro tubes (Sarstedt, cat no. 72.693) and stored at −20 °C.

### Sample preparation

All sample handling, DNA isolation, and library preparations were performed inside a UV hood to minimise contamination. Before starting the work, boxes with sterile plastic consumables and containers with reagents were UV-sterilized for at least 1 h in the closed hood. The DNA extraction, library preparation, and sequencing took place between October 5 and November 13, 2015. Sample slurries were prepared from the five different sample types with guidance from Flores *et al.* (2012). Ileal, caecal, colon, and faecal samples were thawed, kept on ice, and vortexed vigorously. From the ileal and caecal samples, 50-100 μl was taken using a pipette, and from the colon and faecal samples, the tip of a sterile disposable spatula was used. The samples were dissolved in 2 ml of Nuclease-free water (Ambion, cat no. AM9938) in 50 ml sterile tubes. For cloacal and control swab samples, 250 μl of sterile water was added directly to the original tube containing the swab.

The tubes were vigorously vortexed for 30 s and slurries (1 ml of gut and faecal slurries and 100 μl of swab slurries) were immediately transferred to pre-sterilised X30 Deep 96 well plates (Axygen, cat no. P-DW-20-C-S-IND 96) and kept on ice. Plates were sealed with X50 Sealing mat ImpermaMat chemical resistant (Axygen, cat no. AM-2ML-RD-IMP) and kept at −25 °C until DNA isolation. Each sample slurry was used in both the subsequent DNA extraction and the direct PCR method, creating two identical sample sets for comparison of the two preparation methods.

### DNA isolation and 16S library preparation using the DNA extraction method

We used the PowerSoil-htp 96 well soil DNA isolation kit (MoBio Laboratories, cat no. 12955-4). The DNA isolation procedure was according to the manufacturer's protocol with slight modifications (Flores *et al.* 2012) (EMP DNA Extraction Protocol, version 4_13, http://www.earthmicrobiome.org/emp-standard-protocols/16s/) as detailed below. Sample slurry plates were thawed and shaken for 90 s at 20 Hz in a TissueLyser (Qiagen) to mix and then spun down (up to 200 xg) in a plate centrifuge. A volume of 25 μl slurry was pipetted from the slurry plates to MoBio PowerSoil-htp Bead Plates, 60 μl of Solution C1 was added, plates were sealed, and incubated during 10 min at 65 °C before the TissueLyser step in the manufacturer's protocol, which was followed thereafter. DNA was eluted in 100 μl Solution C6.

Amplicon libraries (one for each sample) targeting the V3 and V4 regions of the 16S small-subunit rRNA gene were prepared for sequencing according to the Illumina 16S Metagenomic Sequencing Library Preparation Guide (Part # 15044223 Rev.B) with slight modifications detailed below. Each PCR reaction (25 μl) contained 10 μl DNA-extract, 0.5 μM of each Illumina fusion primer with the gene-specific forward and reverse primers Bakt_341F and Bakt_805R (Herlemann *et al.* 2011) and 1x Phusion High-Fidelity PCR Master Mix with HF Buffer (ThermoScientific, cat no. F-531S). The cycling conditions were: 98 °C for 30 s followed by 25 cycles of 98 °C for 10 s, 56 °C for 15 s and 72 °C for 20 s. A final extension step at 72 °C for 10 min was applied. The amplicons were purified with AmPure XP beads (Agencourt, cat no. A63881) in a ratio of 1:0.8 (PCR-product:bead solution) and freshly prepared 80% ethanol according to the manufacturer's protocol.

The purified amplicons were each dissolved in 43 μl Nuclease-free water (Ambion, cat no. AM9938) and 15 μl was transferred to a new PCR-plate for subsequent dual index PCR with the Nextera Index Kit V2 sets C or D (Illumina, cat no. FC-131-2003 and FC-131-2004) to individually label the amplicons. The index PCR reactions (50 μl) contained 5 μl each of forward and reverse Nextera Index primers and 1x Phusion High-Fidelity PCR Master Mix with HF Buffer (ThermoScientific, cat no. F-531S). The cycling conditions were: 98 °C for 30 s followed by 8 cycles of 98 °C for 10 s, 62 °C for 30 s and 72 °C for 30 s. A final extension step was applied at 72 °C for 10 min.

The individually indexed amplicons were purified with AmPure XP beads (Agencourt, cat no. A63881) in a ratio of 1:1.12 (PCR-product:bead solution) and freshly prepared 80% ethanol according to the manufacturer's protocol. The purified individually indexed amplicons were then each dissolved in 43 μl Nuclease-free water (Ambion, cat no. AM9938) and 38 μl was transferred to a new PCR plate. Amplicons were quantified on the plate reader system FLUOstar Omega (BMG Labtech) using Quant-iT PicoGreen dsDNA Kit (Invitrogen, cat no. P7589). Weak PCR-products (<4.5 ng/μl) were evaporated to increase DNA concentration and then re-quantified. All amplicons were pooled in equimolar amounts and were analysed on a 2100 BioAnalyzer (Agilent Technologies, cat no. G2940CA) before sequencing.

### 16S library preparation using the direct PCR method

The direct PCR approach was performed using the Extract-N-Amp Plant PCR kit (SigmaAldrich, cat no. XNAP2) with guidance from Flores *et al.* (2012). Sample slurry plates were thawed and shaken for 90 s at 20 Hz in a TissueLyser (Qiagen) to mix and then centrifuged (up to 200 xg) in a plate centrifuge. A volume of 25 μl slurry was pipetted from each sample in the slurry plates into 100 μl Extract-N-Amp Plant PCR kit Extraction buffer in pre-sterilised X30 Deep 96 well plates (Axygen, cat no. P-DW-20-C-S-IND 96). Plates were sealed with X50 Sealing mat ImpermaMat chemical resistant (Axygen, cat no. AM-2ML-RD-IMP). The plates were heated in a water bath for 10 min at 95 °C and then centrifuged at 2500 xg for 5 min before adding 100 μl Extract-N-Amp Plant PCR kit Dilution buffer to each well by gentle pipetting. The plates were kept at −25 °C until direct PCR.

Amplicon libraries were created for the resulting direct PCR-extracts. The same Illumina-adaptor modified forward primer Bakt_341F and reverse primer Bakt_805R (Herlemann et al. 2011) as for the DNA extraction method samples were used. Each PCR reaction (25 μl) contained 5 μl Extract-N-Amp DNA extract, 0.5 μM of each Illumina fusion primer, and 1x Extract-N-Amp PCR Ready Mix (SigmaAldrich, cat no. XNAP2). The cycling conditions were: 94 °C for 3 min followed by 25 cycles of 94 °C for 30 s, 50 °C for 30 s, and 72 °C for 45 s. A final extension step at 72 °C for 10 min was applied.

The amplicons were purified, individually labelled, quantified and pooled for sequencing using the same method as described above for the samples prepared with the DNA extraction method. The direct PCR samples were labelled with Nextera Index Kit V2 sets A or B (Illumina, cat no. FC-131-2001 and FC-131-2002).

### Replicate samples

We created two sets of replicate samples for each preparation method in order to evaluate repeatability. The first set, “extraction replicates”, included a total of 80 sample pairs (n = 40 pairs for each method, partitioned into 8 pairs per sample type) and was created before the DNA extraction and the direct PCR procedures, by dispensing the slurry from the same sample into two wells situated on two different plates. The second set, “PCR replicates”, was created by amplifying from the same DNA-extract in two separate PCR wells. The PCR replicates included a total of 20 pairs of samples (n = 10 pairs for each method, partitioned into 2 pairs per sample type). These two replicate levels were created to evaluate at what stage in the library preparation procedure we could detect potential differences, if repeatability differed.

### Amplicon sequencing

DNA sequencing was performed according to the Illumina 16S Metagenomic Sequencing Library Preparation Guide (Part # 15044223 Rev.B) at the DNA Sequencing Facility, Department of Biology, Lund University. The equimolarly pooled amplicon libraries and 10% PhiX spike-in using the PhiX Control Kit V3 (Illumina, cat no. FC-110-3001) were sequenced in one 300-bp paired end run on an Illumina MiSeq platform using the MiSeq Reagent Kit V3 (600 cycles) (Illumina, cat no. MS-102-3003). The pool of amplicon libraries (and PhiX) was added as 8 pM and produced at a cluster density of 891 K/mm2. To summarise, a total of 321 different amplicon libraries were part of this study. The amplicon libraries represented 100 unique ostrich gut microbiome samples (n = 20 per sample type), 2 control and 2 blank samples, 40 extraction replicates, and 10 PCR replicates, that were all prepared both with the DNA extraction and the direct PCR method (Table S1). An additional 11 control and 2 blank samples from a subsequent run were also evaluated to increase the number of controls (Table S1).

### Data processing

The 16S amplicon sequences were quality controlled using FastQC (v. 0.11.5) (Andrews 2010) together with MultiQC (Ewels *et al.* 2016). Primers were removed from the sequences using Trimmomatic (v. 0.35) (Bolger *et al.* 2014) and the forward reads were retained for analyses. Quality filtering of the reads were executed using the script multiple_split_libraries_fastq.py from QIIME (v. 1.9.1) (Caporaso *et al.* 2010). All bases with a Phred score < 25 at the 3’ end of reads were trimmed and samples were multiplexed into a single high-quality multi-fasta file.

Operational taxonomic units (OTUs) were assigned and clustered using Deblur (v. 1.0.0) (Amir *et al.* 2017). Deblur circumvents the problems surrounding clustering of OTUs at an arbitrarily threshold by obtaining single-nucleotide resolution OTUs after correcting for Illumina sequencing errors. This results in exact sequence variants (ESVs), also called amplicon sequence variants (ASVs), oligotypes, and sub-OTU (sOTUs). In order to avoid confusion, we chose to call these units OTUs, but the reader should be aware that they differ from the traditional 97% clustering approach (Callahan *et al.* 2017). The minimum reads-option was set to 0 to disable filtering inside Deblur, and all sequences were trimmed to 220 bp. We used the biom table produced after both positive and negative filtering, which by default removes any reads which contain PhiX or adapter sequences, and only retains sequences matching known 16S sequences. Additionally, PCR-originating chimeras were filtered from reads inside Deblur (Amir *et al.* 2017).

Taxonomic assignment of OTUs was performed using the Greengenes database (DeSantis *et al.* 2006). We filtered all samples on a minimum read count of 1000 sequences, resulting in six out of 321 samples being directly excluded (three blank and three ileal samples). We further filtered all OTUs that only appeared in one sample, resulting in 4,290 OTUs remaining out of an initial 18,689. All samples with technical replicates (both extraction and PCR replicates) had the sequence data merged within their respective sample type (i.e. ileum.rep1 + ileum.rep2) to increase the amount of sequence information per sample for all analyses except repeatability analyses where the replicates were evaluated separately. We present results from non-rarefied data in this study, as recommended by McMurdie & Holmes (2014).

### Data analyses

Analyses were performed in R (v. 3.3.2) (R Core Team 2017), and plots were made using phyloseq (McMurdie & Holmes 2013) and ggplot2 (Wickham 2009). We calculated alpha diversity of samples using the Shannon measure with absolute abundance of reads, and distance measures with the Bray-Curtis distance method on relative read abundances in phyloseq (v. 1.19.1) (McMurdie & Holmes 2013). Community level microbiome differences between the direct PCR and the DNA extraction method were examined using permutational multivariate analysis of variances (PERMANOVA) on Bray-Curtis distances using the Adonis function in vegan (v. 2.4-2) with 1000 permutations (Oksanen *et al.* 2017). Sequencing of blank (negative) samples resulted in extremely few sequence reads: three blank samples had < 320 reads and the other three had < 3,000 reads, compared to an average of 10,689 reads for the other sample types (Figure S1). Control swabs showed highly dissimilar microbial composition to all other samples (see Videvall *et al.* 2017) and therefore we did not include these in any further analyses.

To evaluate bacterial abundances, we first filtered out all OTUs with less than 10 sequence reads and then, using DESeq2 (v. 1.14.1), counts were modelled with a local dispersion model and normalised per sample using the geometric mean (Love *et al.* 2014). Differential abundances between the preparation methods were subsequently tested in DESeq2 with a negative binomial Wald test using individual ID as factor and with the beta prior set to false (Love *et al.* 2014). The results for specific comparisons were extracted (e.g. faeces-DirectPCR versus faeces-Extraction) and p-values were corrected with the Benjamini and Hochberg false discovery rate for multiple testing (Benjamini & Hochberg 1995). OTUs were labelled significant if they had a corrected p-value (q-value) < 0.01.

We examined the repeatability of the two methods by evaluating the strength of the correlation in normalised OTU abundance between paired sample replicates. This was done separately for the two methods, and for the two replicate sets (extraction replicates and PCR replicates). Correlation coefficients were calculated using Spearman’s rank correlations on all OTUs with non-zero abundances.

## Results

### Practical aspects of direct PCR and DNA extraction

The total time spent extracting DNA using the direct PCR method was considerably shorter (45 minutes) compared to the conventional DNA extraction method (8 hours) (Table 1). The cost of using the direct PCR method was also lower, as were the number of steps in each protocol (Table 1). Nevertheless, the number of sequence reads obtained per sample (mean = 10,689) did not differ between the DNA extraction method and the direct PCR method (two sample t-test: t = 1.25, df = 290.7, p = 0.21) (Figure S1), as expected since equimolar PCR products from the samples were combined before sequencing.

### Description of the microbiomes obtained with direct PCR and DNA extraction

Samples clustered strongly according to sample type both in the Principal Coordinates Analysis (PCoA) (Figure 1A) and the network analysis (Figure 1B), although some minor separation between the direct PCR and DNA extraction methods was evident. The two library preparation methods yielded fairly consistent patterns for the total number of OTUs per bacterial class and sample type, but also reflected some notable differences in taxa composition (Figure 1C). Specifically, Bacilli were slightly more abundant in all sample types with the DNA extraction method relative the direct PCR method, and the ileum in particular showed the largest class differences, with a higher abundance of Mollicutes, Gammaproteobacteria, and Bacteroidia with the direct PCR method (Figure 1C).

**Figure 1.**
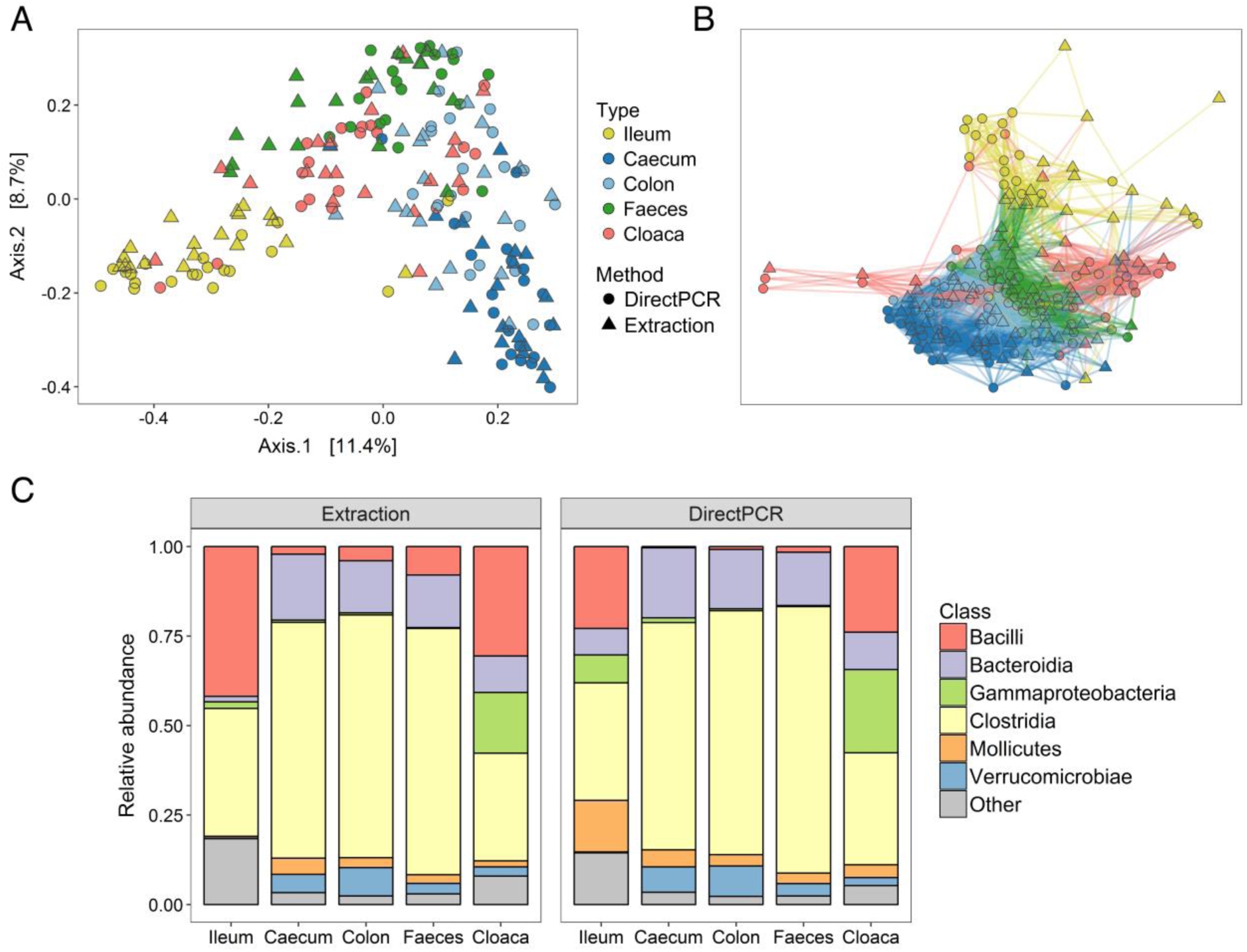
Microbiomes of different gut samples obtained using direct PCR and conventional DNA extraction methods. (A) Principal coordinates analysis (PCoA) and (B) network analysis of all sample types from all individuals prepared using the two methods. (C) Relative abundances of bacterial classes in the different sample types from the two methods.

To further investigate if the microbiota of samples from the direct PCR and DNA extraction methods differed depending on gut site, we performed separate PCoAs for each of the five sample types. The caecum, colon, and faeces showed very high correspondence in beta diversity for identical samples prepared using direct PCR and DNA extraction, as they clustered by individual and not method, whereas differences were much greater for cloacal and ileal samples (Figure 2).

**Figure 2.**
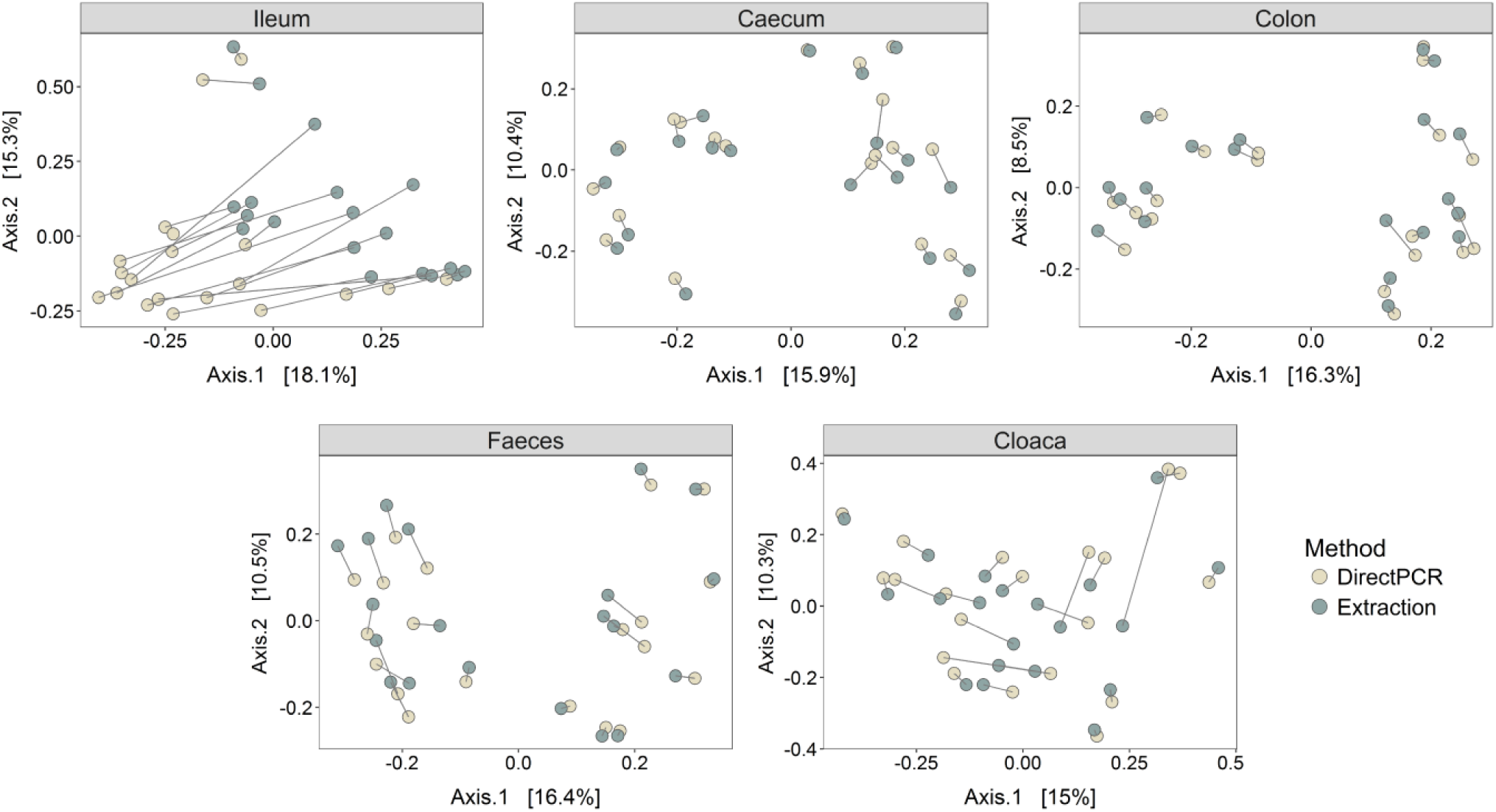
Principal coordinates analyses (PCoAs) of Bray-Curtis distances of different sample types. Lines between points denote identical samples prepared using direct PCR (light dots) and DNA extraction (dark dots) methods. Numbers within brackets show percent variance explained by the PCoAs.

### Differences in microbiomes obtained with direct PCR and DNA extraction

We next evaluated differences in alpha diversity (OTU richness) between direct PCR and DNA extraction methods for the different sample types (Figure 3A). There were no differences in alpha diversity between the two methods in the colon, faeces, and cloaca (paired Wilcoxon signed rank test: n_pairs_ = 20 per type, p > 0.4) (Figure 3A). However, alpha diversity was significantly higher in the direct PCR samples compared to DNA extraction samples for the ileum (n_pairs_ = 19, V = 5, p < 0.0001), and significantly lower in the direct PCR caecal samples (n_pairs_ = 20, V = 199, p = 0.0001) (Figure 3A). Correlation analyses of alpha diversity between the two methods also showed higher diversity for ileal direct PCR samples and slightly lower diversity for caecal direct PCR samples, but the strength of the correlations between methods were generally high for all sample types (r = 0.56–0.91) (Figure S2).

**Figure 3.**
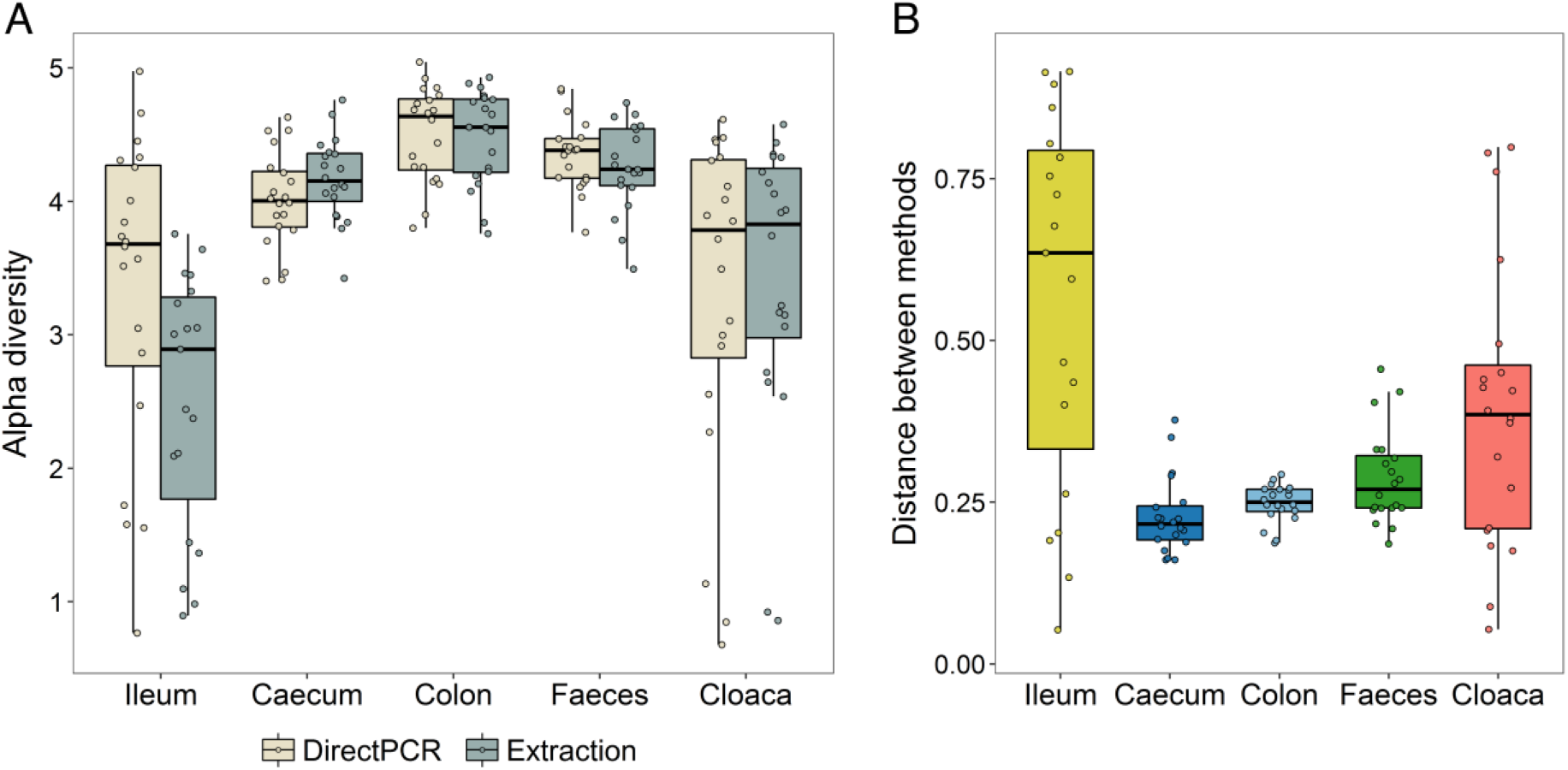
Differences in the microbiomes obtained using direct PCR and DNA extraction methods. (A) Alpha diversity (Shannon index) of all samples within the different sample types, for the direct PCR and the DNA extraction methods. (B) Bray-Curtis distances between identical samples, prepared with direct PCR and DNA extraction methods.

Dissimilarities in microbiome composition between samples, as calculated by the Bray-Curtis distance measure, showed significant effects of method, sample type, individual, and the interaction between method and sample type (PERMANOVA: all effects: p < 0.001). The overall variance explained by method and method*sample type was, however, extremely small (R2 = 0.014, and R2 = 0.019, respectively), whereas the variance explained by host individual (R2 = 0.283) and sample type (R2 = 0.201) were substantially larger.

Examining the Bray-Curtis distances between the two methods within sample types revealed that while the caecum, colon, and faeces all showed relatively low distances (mean: 0.23, 0.25, and 0.29, respectively), the cloaca (mean = 0.39) and most notably, the ileum (mean = 0.56), displayed much greater distances and much higher variances (Figure 3B). Specifically, the distances between identical samples prepared with each method from the ileum were significantly higher than the corresponding distances in the caecum, colon, and faeces (n_pairs_ = 19, V >= 165, q < 0.009 for each of these three tests) (all pairwise comparisons between sample types [10 tests] tested using paired Wilcoxon signed rank test with p-values corrected for multiple testing). The cloacal samples showed significantly higher distances than caecal (n_pairs_ = 20, V = 21, q = 0.005) and colon samples (n_pairs_ = 20, V = 34, q = 0.013), and finally, the caecal samples showed slightly lower distances than faecal samples (n_pairs_ = 20, V = 40, q = 0.023) (Figure 3B).

### Differences in OTU abundances with direct PCR and DNA extraction

Calculating the correlation of average OTU abundances between the direct PCR and extraction methods revealed very high correlation coefficients for the caecum, colon, and faeces (r_s_ = 0.84–0.86), weaker for the cloaca (r_s_ = 0.60), and negative for the ileal samples (r_s_ = −0.17) (Figure S3).

Analyses of differences in the abundance of specific OTUs between the two methods resulted in very few significantly different OTUs in the caecum (n = 9), colon (n = 13), and faeces (n = 24) (Figure 4). However, there were many more in the cloaca (n = 67), and the ileum demonstrated a staggering 324 significant OTUs between the DNA extraction and the direct PCR method (Figure 4). Notably, the vast majority of significant OTUs across all sample types (80%) had higher abundances when using direct PCR compared to DNA extraction.

**Figure 4.**
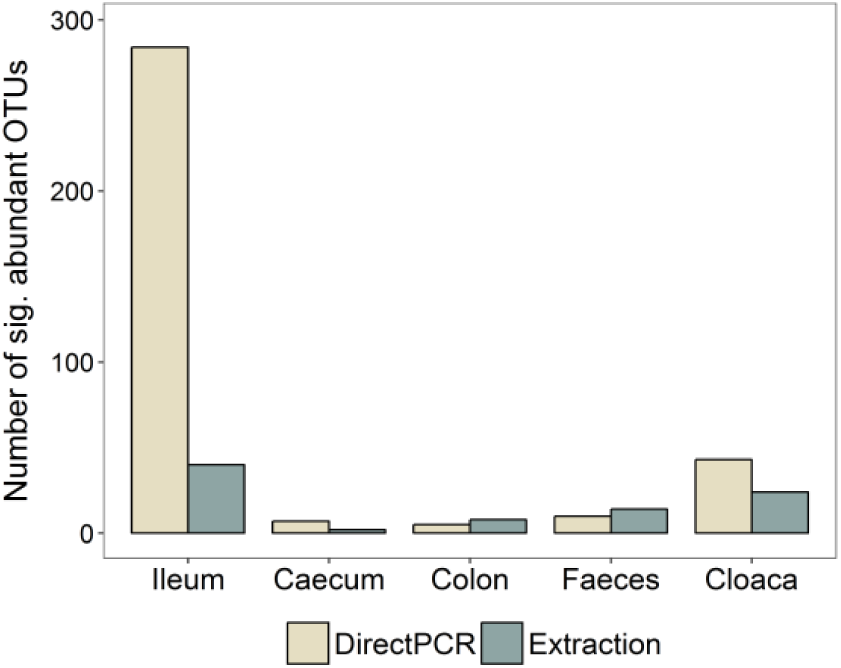
Number of OTUs with significantly higher abundance in either library preparation method for the different sample types.

Comparing the exact OTUs that had significantly different abundances in the five sample types, showed that they were unique to each sample type (i.e. OTUs were only significantly different within one type) (Table S2). We found one genus, however, *Mycoplasma*, with significant OTUs present in all sample types. All significant *Mycoplasma* OTUs (family: Mycoplasmataceae) had higher relative abundances in the samples from the direct PCR compared to the DNA extraction method (Figure 5). Other genera with significantly different abundances in multiple sample types were e.g. *Anaerofustis* (higher abundance with the direct PCR method in colon, faecal, and cloacal samples) and *Klebsiella* (more numerous in the direct PCR method of ileum, caecum, and colon). The genus *Prevotella* (class: Bacteroidia) was the most prevalent in the list of significant genera, representing 43 unique OTUs in the cloaca and ileum, all of which (100%) had higher abundance in the direct PCR samples (Table S2).

**Figure 5.**
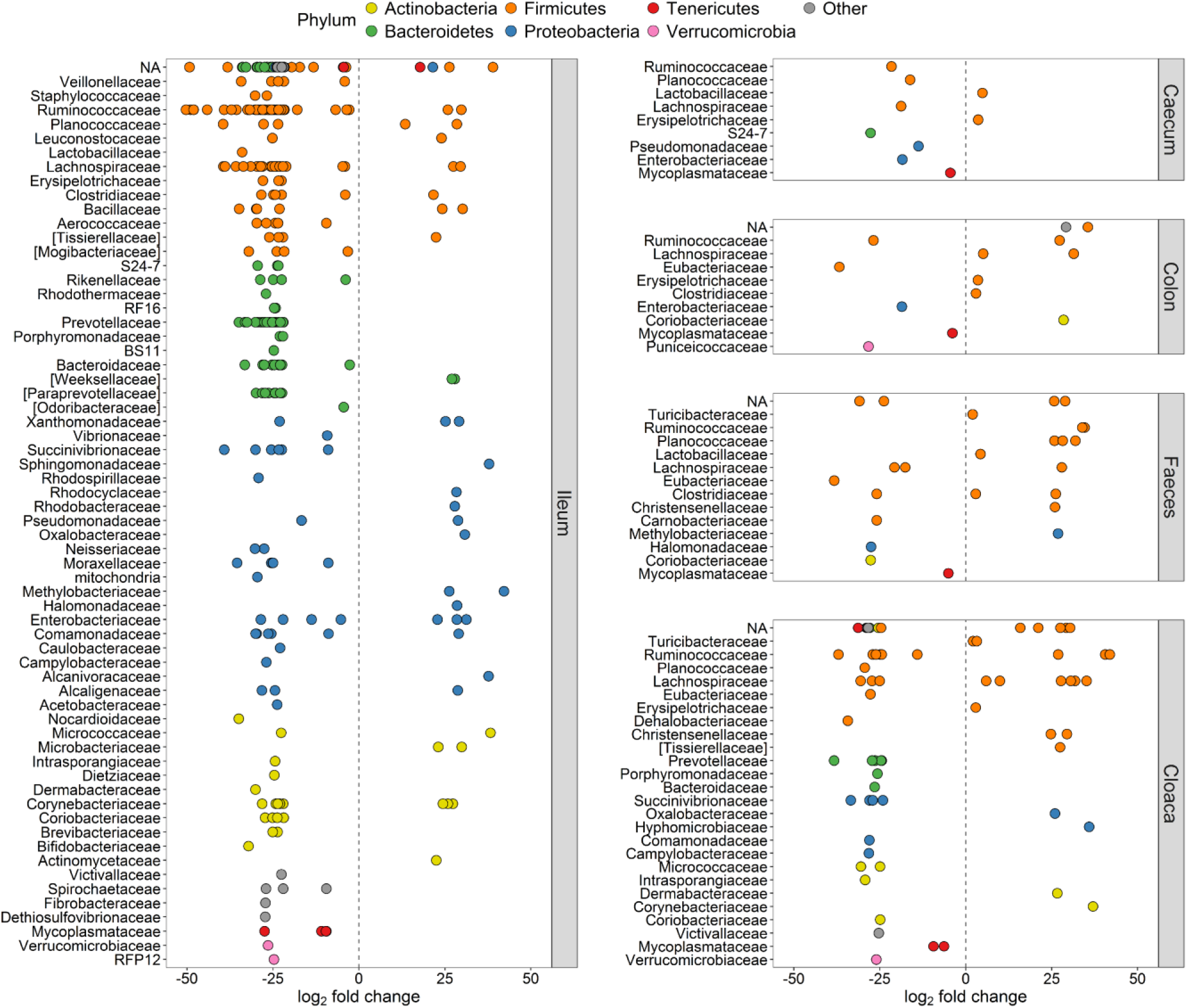
Significantly differentially abundant OTUs (q < 0.01) between direct PCR and conventional DNA extraction methods. Y-axes show taxonomic families and all OTUs have been coloured within their respective phylum and separated per sample type. Positive log_2_ fold changes indicate higher relative OTU abundance in the DNA extraction method, and negative log_2_ fold changes signify higher abundance in the direct PCR method.

The phylum with the highest number of significantly differentially abundant OTUs was the Firmicutes (n = 210 in total), and in particular the class Clostridia (n = 171 in total) (Figure 5; Table S2), which was in majority in most sample types (Figure 1C). The genera which comprised the most significant differentially abundant OTUs between the two methods were the *Oscillospira* (class: Clostridia), *Mycoplasma* (class: Mollicutes), and *Coprococcus* (class: Clostridia) (Table S2).

### Repeatability of replicate samples with direct PCR and DNA extraction

Next, we evaluated the repeatability of the DNA extraction and direct PCR methods by calculating correlations of OTU abundances and diversity between pairs of replicate samples. For the “extraction replicates”, the correlation coefficient of OTU abundance was almost identical for the DNA extraction method (r_s_ = 0.73; Figure 6A) and the direct PCR method (r_s_ = 0.70; Fig 6B). For the “PCR replicates”, the strength of the correlation was slightly higher but again similar for the two methods (DNA extraction: r_s_ = 0.82, Figure 6C; and direct PCR: r_s_ = 0.80, Figure 6D).

**Figure 6.**
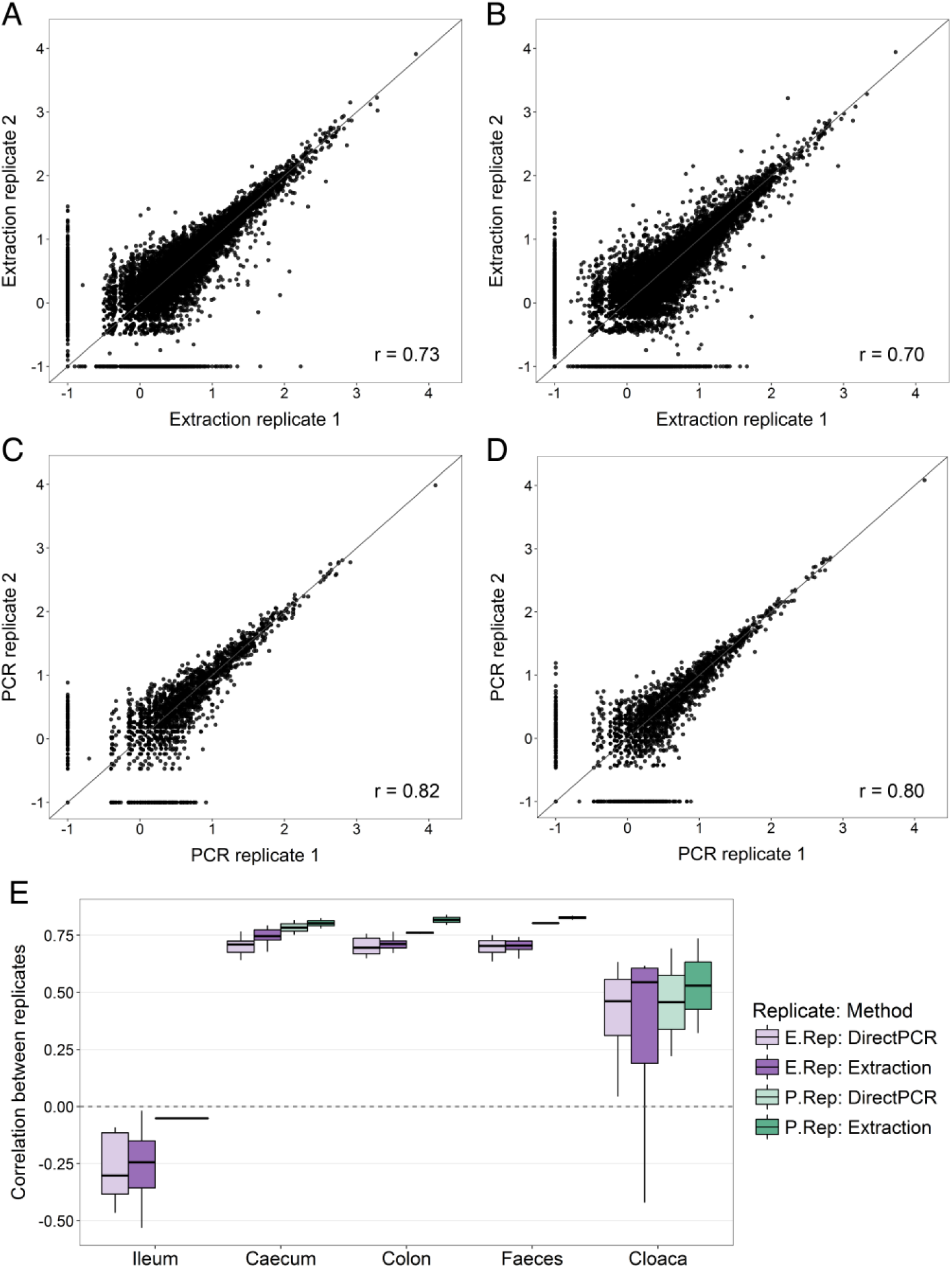
Repeatability of OTU abundances between replicate samples. Scatter plots show abundances of all OTUs in (A) extraction replicates obtained by the DNA extraction method, (B) extraction replicates obtained by the direct PCR method, (C) PCR replicates obtained by the DNA extraction method, and (D) PCR replicates from the direct PCR method. OTU abundances have been normalised according to the DESeq2 method and log-transformed (+0.1) for graphical purposes. r values show the Spearman’s rank correlation coefficients. (E) Boxplot of correlation coefficients (Spearman’s rank correlation) of OTU abundances calculated separately for all 100 replicate sample pairs.

When we partitioned the OTU abundance data according to sample type, large differences in repeatability were observed (Figure 6E). The caecal, colon, and faecal samples had the strongest correlations between replicates, with both methods having an average r_s_ = 0.70–0.74 for the extraction replicates and an average r_s_ = 0.76–0.83 for the PCR replicates (Figure 6E). In contrast, the extraction replicates from the cloaca were characterised by a much weaker correlation (r_s_ = 0.36), as did the cloacal PCR replicates (r_s_ = 0.49), and the correlations between ileal replicates were even negative (extraction replicates: r_s_ = −0.27, PCR replicates: r_s_ = −0.05) (Figure 6E).

Finally, we examined the correlation between alpha diversity estimates in the replicate samples to evaluate the repeatability of the community between methods. Relative to the OTU abundance data, there was higher repeatability in alpha diversity for cloacal and ileal samples (r = 0.79–0.96), while the caecal, colon, and faecal samples again had high repeatability for both methods (r = 0.70–0.97) (Figures S4–S5).

## Discussion

This study shows that direct PCR provides highly comparable results to the widely used and recommended DNA extraction method in analyses of gut microbiomes of animals. Both techniques give qualitatively and quantitatively similar estimates of microbial diversity and abundance for caecal, colon, and faecal samples, and were highly repeatable for these sample types. However, the two methods present dissimilar microbiomes for cloacal, and in particular, ileal samples, recovering large differences and poor repeatability in OTU abundances across replicates. We discuss hypotheses that may explain why these methods perform well with some sample types, but not others.

During amplicon library preparation, PCR products from the ileal and cloacal samples were much weaker relative to the other sample types, presumably due to lower microbial biomass in these samples. Low initial DNA template concentration may reduce repeatability in two ways. First, low DNA concentrations will introduce stochasticity in what bacterial species in the original samples are amplified. Second, small differences between the direct PCR and DNA extraction methods can become exaggerated in low DNA samples because the sequence coverage will be greater for low abundance OTUs as a result of equal quantities of PCR product being used from all samples before sequencing. However, the uniform direction observed between identical samples in the PCoA of ileum (Figure 2) indicated that there were consistent discrepancies between the two methods. This was also supported by the correspondence in alpha diversity of cloacal and ileal samples both between replicates (Figures S4−S5) and between the methods (Figure S2). This suggests that although it is difficult to measure relative abundances of specific bacterial taxa when DNA concentrations are low, it may still be possible to gain an accurate measure of the community composition using both of these methods.

The direct PCR samples from the ileum had significantly higher alpha diversity (Figure 3A) and higher relative abundances in the majority of differentially abundant OTUs (Figure 4), compared to the DNA extraction method. One potential reason for this difference is that more DNA is lost during the DNA extraction procedure, which is column-based with several wash and transfer steps. This is known to be associated with high DNA loss, whereas with direct PCR, the individual samples are contained in just one plate well during the full extraction procedure. In samples with low starting DNA concentrations, direct PCR may therefore be superior to conventional extraction methods at recovering rare bacterial taxa. The higher diversity in the ileal direct PCR samples could also be a consequence of the regular Taq polymerase included in the direct PCR reaction mix, which is slightly more error-prone than the Phusion High-Fidelity polymerase used for amplification in the DNA extraction protocol. A higher error rate during the direct PCR amplification step could potentially yield spurious OTUs, raise the alpha diversity and the number of differentially abundant OTUs. However, Taq polymerase does not explain the consistent changes observed in the PCoA (Figure 2), the correlation in diversity between the methods (Figure S2), or the changes in relative abundances of specific taxonomic groups (Figure 1C and Figure 5).

Despite high correspondence in abundance estimates across most OTUs, there were some consistent differences in the abundance of specific taxa between direct PCR and the DNA extraction method. It is possible that morphological differences between bacterial groups may influence the efficiency with which the two methods recover DNA. For example, compared to the DNA extraction method, we found that direct PCR had higher relative abundances of Mollicutes, which are small bacteria without cell wall, and fewer Bacilli, gram-positive bacteria with a thick cell wall, in the ileal samples. The direct PCR method uses chemical and heat shock treatment to lyse cells, whereas the DNA extraction method lyses cells with mild heat, chemical, and mechanical treatment. This could in theory have different effects on the release of DNA according to the bacterial cell wall structure. The direct PCR approach previously tested on human samples (Flores *et al.* 2012) found Prevotellaceae to be in higher abundance in several tongue samples and Veillonellaceae more abundant in gut samples. We obtained similar results, with 43 significant OTUs belonging to Prevotellaceae, and five significant Veillonellaceae, all more abundant in the ileal and cloacal samples from the direct PCR method (Figure 5), suggesting that direct PCR may be better at recovering these specific taxa. Furthermore, we found *Mycoplasma* spp. present in significantly higher abundance in all sample types with the direct PCR method (Table S2). *Mycoplasma* are sometimes present as laboratory contaminants (Drexler & Uphoff 2002), however sequencing negative controls of the direct PCR kits produced practically zero occurrences of *Mycoplasma* (0−2 reads), and extremely few sequences of other bacteria (66−193 reads), proving contamination by the direct PCR kits highly unlikely.

In summary, direct PCR and conventional DNA extraction methods gave highly similar estimates of community composition and overall OTU abundance in gut microbiomes from high microbial biomass samples, such as those from the caecum, colon, and faeces. The direct PCR protocol is, however, cheaper, takes less time, and involves 29 fewer laboratory steps, thereby reducing the risk of contamination and human errors (Table 1). These practical advantages, combined with the fact that direct PCR is possibly more sensitive at measuring diversity in low DNA concentration samples, highlight direct PCR as an excellent substitute for DNA extraction methods. We hope that these results will aid researchers in the planning of gut microbiome studies on animals.

## Acknowledgements

We are grateful to Adriaan Olivier, Maud Bonato, Naomi Serfontein, Julian Melgar, and all staff at the Oudtshoorn Research Farm, Western Cape Government for assisting with sample collection. Funding was provided partially by the Helge Ax:son Johnson Foundation, the LÄngmanska Cultural Foundation, the Lund Animal Protection Foundation, the Lars Hierta Memorial Foundation, and the Royal Physiographic Society of Lund to E.V., and by a Wallenberg Academy Fellowship and Swedish Research Council (VR) grant to C.K.C., and by the Western Cape Government.

## Author contributions

M.S., E.V., and C.K.C. planned the study. S.C. provided animal facilities, and A.E. supervised the experimental part of the study. M.S. planned and performed the laboratory work. E.V. performed the data analyses. E.V., M.S., and C.K.C. wrote the paper with input from S.C. and A.E.

## Data accessibility

Supporting information has been made available online. Sequence data have been uploaded to the European Nucleotide Archive (ENA) under accession number: PRJEB22648.

## Supporting information

Supplementary figures have been made available online with the preprint. Supplementary tables are available upon request.

**Table S1.** Sample information.

**Table S2.** Significantly differentially abundant OTUs between methods.

**Figure S1**. Number of reads per sample.

**Figure S2**. Correlation of alpha diversity between methods.

**Figure S3**. Correlation of OTU abundances between methods.

**Figure S4**. Repeatability of alpha diversity between extraction replicates using the DNA extraction method.

**Figure S5**. Repeatability of alpha diversity between extraction replicates using the direct PCR method.

